# The impact of genetic background and gender on the increase in mitotic index in response to mating of *Drosophila melanogaster*

**DOI:** 10.1101/2020.05.15.098509

**Authors:** Manashree Malpe, Cordula Schulz

## Abstract

The replenishment of specialized cells depends on the activity of stem cells. Recent advances in stem cell research have shown that the germline stem cells (GSCs) in *Drosophila melanogaster* can increase their mitotic activity in response to mating. Here, we show that this ability to respond to mating is eliminated if the males are mutant for the ABC transporter, White, the genetic background for a plethora of fly lines. Furthermore, we were not able to reproduce previous findings that female flies increase their GSC numbers and mitotic activity upon mating. Our findings underline the importance of careful experimental design and control specimen.

## INTRODUCTION

As specialized cells are often lost due to usage or death, many tissues have to replenish these through a process commonly referred to as tissue homeostasis. One of the key factors in tissue homeostasis is the ability of long-term precursor cells, the stem cells, to self-renew and to generate short-lived precursor cells that, in turn, undergo a cascade of regulated proliferation and differentiation steps. Stem cells have to adjust their activity to situations of demand. Recent advances in the literature suggests that GSCs in the gonad of *Drosophila melanogaster* can adjust their mitotic divisions to a variety of factors, including diet, temperature, and mating status (Drummond-Barbosa and Spradling, 2001; Parrott et al., 2012; Malpe et al., 2020). For example, when female flies are kept on a yeast-rich diet, their GSCs cycle significantly faster compared to the GSCs of their siblings kept on a yeast-poor diet (Drummond-Barbosa and Spradling, 2001).

Genetic studies in *Drosophila* are highly facilitated by the ability to generate mutants and transgenes for any gene of interest. *Drosophila* transgenes carry morphological markers, such as genes for eye or body color. When transgenes are incorporated into flies lacking the wild-type (wt) versions of these markers, the restoration of the morphological traits allows for easy identification of the transgenes within genetic crosses (Klemenz et al., 1987; Rubin and Spradling, 1982; St. Johnston, 2013). The same morphological traits are often used to mark a mutant chromosome and flies carrying these traits normally serve as genetic backgrounds to study the effect of a mutation on the anatomy, or behavior of the fly (Greenspan, 1997)

One such morphological marker is a mutation in the *white* (*w*) gene, which results in white-eyed instead of red-eyed flies. The *w* gene encodes an ABC transporter involved in the transfer of small molecules across membranes (Mount, 1987). Animals mutant for *w* display an array of abnormalities in addition to the lack of red eye pigment. For example, they are vision impaired, display memory defects, and have abnormal levels of amines in their brains (Diegelmann et al., 2006; Borycz et al., 2008; Krstic et al, 2013).

We previously showed that males increase the mitotic activity of their GSCs in response to mating. This activity was measured as the number of GSCs in mitosis divided by the total number of GSCs, or the GSC M-phase index (MI^GSC^; Malpe et al., 2020). Here, we show that males homozygous mutant for either of two commonly used alleles of *w*, had the same MI^GSC^ as their non-mated siblings, while males from two wt strains, *Oregon R* (*OR*) and *Canton S* (*CS*), increased their MI^GSC^ when repeatedly mated.

When fruit-flies mate, the males transfers seminal fluid together with the sperm to the female reproductive tract, where both get stored in the spermatothecae and the seminal receptacle (Bloch Qazi et al., 2003). Within the seminal fluid are hormones that modify egg laying and reproductive behavior of the females. As a consequence, subsequent mating of female flies is significantly reduced until the hormones have been depleted from their bodies (Chapman et al., 2003). It was previously reported that this low number of female mating events is sufficient to significantly increase GSC numbers and MI^GSC^. Molecularly, this increase in GSC numbers and activity was associated with signaling via the Ecdysone and the Neuropeptide F receptors, and with Sex Peptide (SP; Ameku and Niwa, 2016; Ameku et al., 2018).

When we mated *Drosophila* males from several genetic backgrounds, we did not detect differences in GSC numbers. Furthermore, when we exposed males to only one to three virgin females, we did not detect an increase in MI^GSC^. Instead, we demonstrated that males had to interact with more than three virgin females and for more than 24 hours for an increase in MI^GSC^ to be statistically relevant (Malpe et al., 2020). To compare the effect of mating on the genders, we performed female mating experiments. We learned that, in our laboratory, non-mated and mated wt and *w* mutant females did not show a difference in GSC numbers or MI^GSC^.

## RESULTS and DISCUSSION

### Animals mutant for w did not increase their MI^GSC^ in response to mating

In *Drosophila* males, GSCs are found at the tip of the testes where they attach to somatic hub cells (Figure 1A, 1A’). A GSC division normally results in a new GSC that remains in contact with the hub and a gonialblast that initiates a proliferation and differentiation program that results in the formation of 64 spermatids (Hardy et al., 1979; Fuller et al., 1993; Figure 1A). MI^GSC^ can be investigated by employing an immuno-fluorescence protocol, in which the hub cells are labeled with an antibody against FasciclinIII (FasIII) and the GSCs with an antibody against Vasa. The Vasa-positive GSCs next to the hub can then be imaged (Figure 1A’) and counted in all focal planes. By adding an antibody against phosphorylated Histone-H3 (pHH3) cells in mitosis become apparent and the MI^GSC^ can be calculated (Parrott et al., 2012).

**Figure 1.**
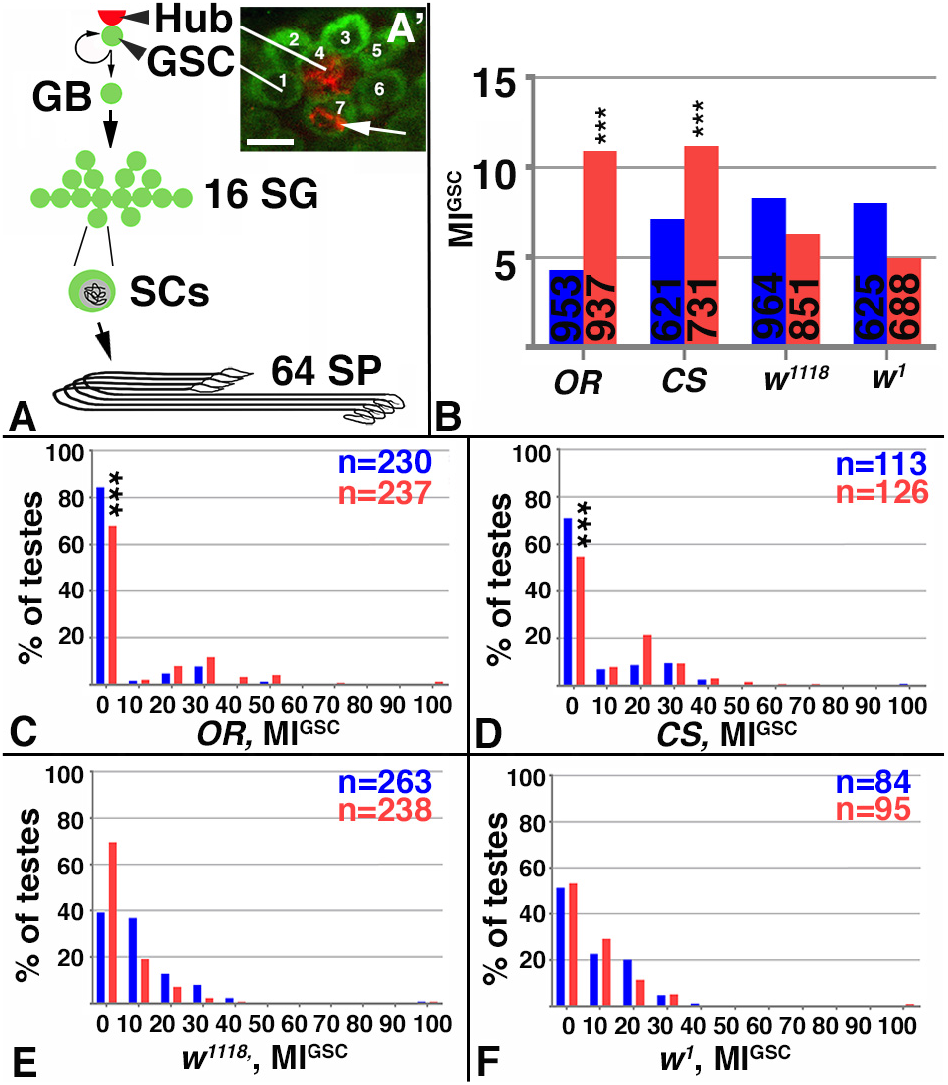
Mated *w* mutant males failed to increase MI^GSC^. A) Cartoon depicting how a GSC division results in a new GSC and a stem cell daughter that will ultimately produce 64 spermatids (SPs). GB: gonialblast, SGs: spermatogonia, SC: spermatocyte. A’) Immuno-staining to the apical tip of a *wt* testis, showing seven Vasa-positive GSCs (green) next to the FasIII-positive hub. One of the seven GSCs is in mitosis based on anti-pHH3-staining (arrow). Scale bar: 10μm. B-F) Blue: non-mated condition, red: mated condition, ***: P-value < 0.001, numbers of GSCs and number of gonads (n=) as indicated. B) Bar graph showing MI^GSC^ of wt and *w* mutants, as indicted. C-F) FDGs showing median of bin of MI^GSC^ across populations of males on the X-axes (bin width=10) and the percentage of testes with each MI^GSC^ on the Y-axes; C) *OR*, D) *CS*, E) *w^1118^*, and F) *w^1^* males.

While wt males always significantly increase MI^GSC^ in response to repeated mating, we noted considerable inconsistencies in our data when males were kept in a *w* mutant background. Intrigued by this observation, we set out to investigate if the failure of males to increase MI^GSC^ might be due to the *w* mutation itself. We obtained two of the commonly used *w* alleles, *w^1118^* and *w^1^*, and set up mating experiments with these, as well as *OR* and *CS* males. All four groups of males mated with the virgin females based on visual observations and the production of progeny (Table 1). However, while *OR* and *CS* males clearly increased MI^GSC^ when mated, animals mutant for *w^1118^* or *w^1^* displayed similar MI^GSC^ under mated and non-mated conditions (Figure 1B).

**Table 1.** Fertility Assay. Male fertility was calculated based on the % of females that produced offspring after mating with males of the indicated genotype. BL#: Bloomington stock number.

| Genotype | BL # | Male fertility |
| --- | --- | --- |
| *OR* | N/A | 72% |
| *CS* | N/A | 62% |
| $w^{1118}$ | N/A | 81% |
| $w^1$ | 679 | 70% |

Figure 1B shows bar graphs, in which the MIs^GSC^ are displayed on the y-axis, while the genotypes are displayed on the x-axis. Bar graphs summarize data across a whole population. In our case, each single bar shows the total number of pHH3-positive GSCs among the total number of GSCs from all testes within one group of males. The frequency distribution graphs (FDGs) in Figure 1C-F, on the other hand, split the same data into smaller units. They show how often different MIs^GSC^ are represented among the testes of one group of males, and, thus, present the data in a less processed way. The FDGs (Figure 1C-F) demonstrate that most testes did not have any GSCs in division (MI^GSC^ of zero), suggesting that GSC divisions are fairly rare. The remainders of the testes normally showed one or more GSCs in division. For both wt strains, the numbers of testes without any GSCs in divisions were significantly reduced upon mating and the remainder of the testes had higher MI^GSC^ compared to non-mated males (Figure 1C,D). For the *w* mutant animals, in contrast, the number of testes with a MI^GSC^ of zero was not reduced upon mating and the remaining testes had about the same MIs^GSC^ as the testes of their non-mated siblings. We conclude that GSCs in *w* mutant animals do not reflect the behavior of wt GSCs with respect to increasing their mitotic activity upon mating.

The *w* mutation is heavily used as a marker in genetic and molecular studies, but the underlying cellular and biochemical functions of *w* are normally not considered as contributing factors to a phenotype. Interestingly, almost all transgenic constructs with the *w* or mini-*w* genes appear to carry an insulator sequence that could have impacted the outcomes of many studies (Chetverina, 2008). Besides the presence of this insulator in transgenic constructs, a variety of abnormal behaviors has been associated with the *w* mutation. For example, *w* mutant flies have altered sensitivity to anesthetic treatments and alcohol, and are less aggressive than wt flies (Campbell and Nash, 2001; Hoyer et al., 2008; Chan et al., 2014). Notably, lack or misexpression of *w* affects sexual behavior in male but not female flies. Expression of *w* throughout the flies from a heat-shock driven transgene, or loss of *w* function caused increased male sexual arousal and male to male courtship behavior, even in the presence of females (Zhang and Odenwald, 1995, Hing and Carlson, 1996; Anaka et al., 2008; Krstic et al., 2013).

The effects of the *w* mutation outside the eye is likely due to a role for *w* in the nervous system as previous studies have shown that *w* mutant animals have abnormal levels of amines, including dopamine and serotonin (Borycz et al., 2008; Anaka et al., 2008; Hoyer et al., 2008). Possibly, the altered amine levels in *w* mutants are due to defects in intra-cellular transport. Animals mutant for *w* have abnormal amine vesicle populations and contents, and display neurodegenerative defects (Borycz et al., 2008; Ferreiro et al., 2017). Consistent with a role for *w* in intra-cellular transport, the W protein was detected in vesicular fractions of several cell types. Electron-microscopy experiments revealed a localization of W in the membranes of the pigment granules within pigment cells and retinula cells, suggesting that it transports molecules from the cytoplasm to the pigment granules (Mackenzie et al., 2000). In malphigian tubule cells, W was detected on vesicles, and in cultured cells, W was detected on the endosomal compartment (Anaka et al., 2008; Evans et al., 2008). In addition to the multitude of effect caused by the *w* mutation, a range of substrates transported by it have been identified. These include Guanosine 3’-5’ cyclic monophosphate (cGMP), Tryptophan, Kynurenine, Guanine, and Riboflavin (Sullivan et al., 1973; Sullivan and Sullivan, 1975; Howells et al., 1977; Sullivan et al., 1979; Sullivan et al., 1980; Evans et al., 2008). The pleiotrophic effect of *w* on the flies and the variety of substances transported by it suggests that the failure of the mutant males to increase their MI^GSC^ could have multiple underlying reasons, ranging from abnormal mating behavior to metabolic effects.

### Mating did not affect female MI^GSC^ or GSC numbers

According to a previous report, *Drosophila* females significantly increase both, their numbers of GSCs and MI^GSC^ upon mating (Ameku and Niwa, 2016). Just as the sperm, eggs are produced from germline stem cells that self-renew and produce daughter cells that proliferate and differentiate (Figure 2A). These GSCs lie at the tip of the ovary within the germarium (Spradling, 1993). Similar to the male GSCs, the female GSCs are attached to somatic cells and can be identified based on their position using molecular markers: the germline marker, Vasa, the mitosis marker, anti-pHH3, and anti-LaminC, which is expressed at high levels in the female stem cell niche (Figure 2B).

**Figure 2.**
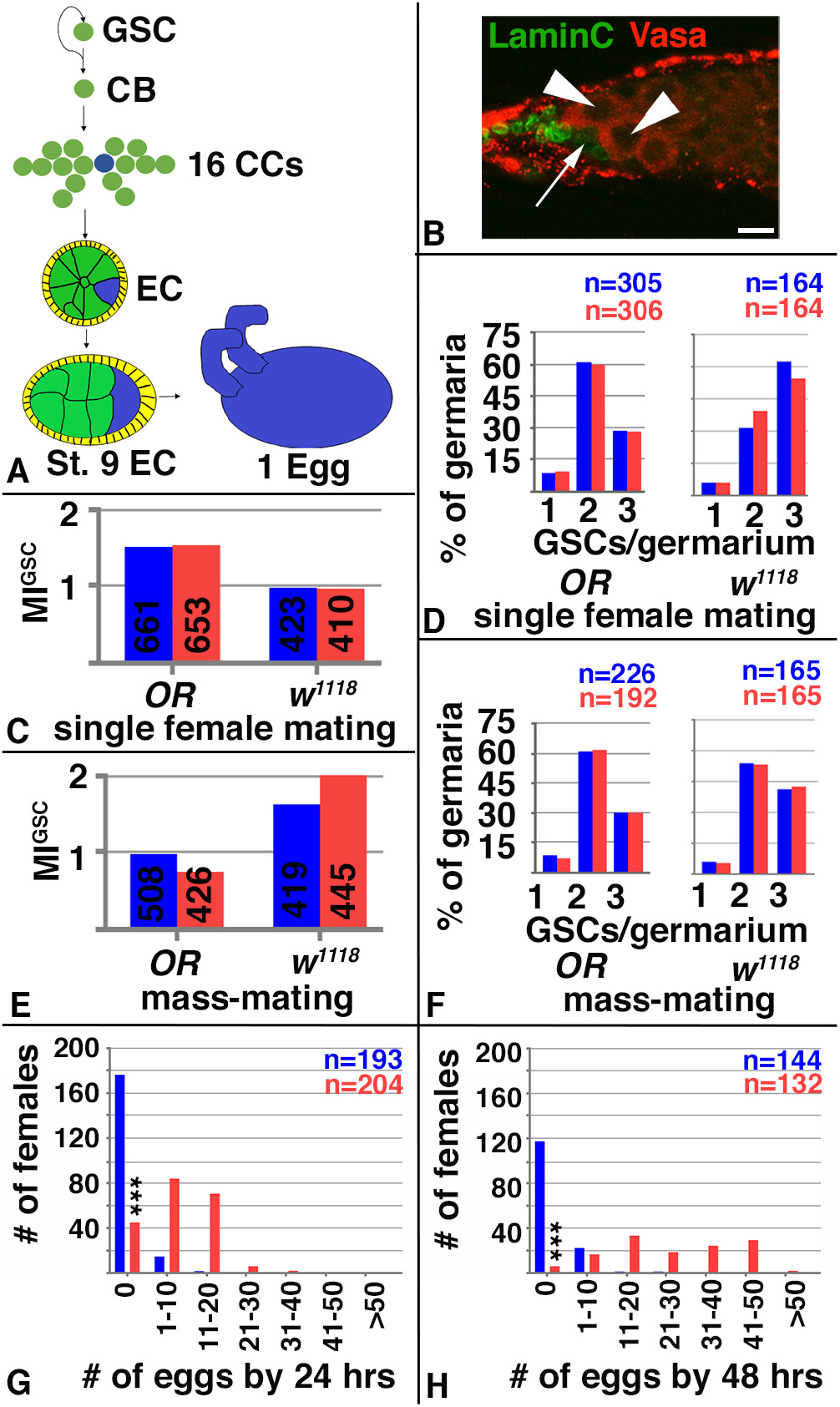
Mating did not affect female MI^GSC^ or GSC numbers but increased oviposition. A) Cartoon illustrating how a female GSC division results in one new GSC and a stem cell daughter that will ultimately produce one egg. CB: cystoblast, CCs: cystocytes, St: stage, EC: egg chamber. B) The apical tip of an *OR* germarium showing two Vasa-positive GSCs (arrowheads) next to LaminC-positive cap cells (arrow). C-H) Blue: virgin condition, red: mated condition, ***: P-value < 0.001, numbers of GSCs and number of gonads (n=) as indicated, genotypes as indicated. C) Bar graph showing no significant difference in MI^GSC^ of virgin and mated females using a single-female mating protocol. No significant difference in GSC numbers was seen between virgin and mated females using a single-female mating protocol. Bar graph showing no significant difference in MI^GSC^ of single virgin and mated females, using a mass-mating protocol. D) No significant difference in GSC numbers was seen between virgin and mated females, using a mass-mating protocol. E) Bar graph showing no significant difference in MI^GSC^ of single virgin and mated females, using a mass-mating protocol. F) No significant difference in GSC numbers was seen between virgin and mated females, using a mass-mating protocol. G, H) FDG showing the numbers of eggs laid by virgin and mated females after G) 24 and H) 48 hours (hrs) of the experiment on the X-axes and the number of females laying the amounts of eggs on the Y-axes.

In a first attempt to reproduce the published data, we mated *OR* and *w^1118^* females to *OR* males for 24 hours in single-female mating experiments. We did not observe an increase in GSC numbers or MI^GSC^ in mated female flies. *Drosophila* females have far more stem cells compared to males. In the males, each of the two testicular tubules has an average of 5-12 GSCs (Hardy et al., 1979; Malpe et al., 2020). Each of the two female ovaries contains 15-20 tubes, called ovarioles, in which the eggs develop. Each of these ovarioles has a germarium at the tip that contains on average two GSCs (Schüpbach et al., 1993; Wieschaus and Szabad, 1979). Thus, one female should have about 30-40 GSCs per ovary. For representative results across the population of females, we only imaged on average 10 germaria from every ovary. For both fly lines, the non-mated and mated females had the same MI^GSC^ (Figure 2C). We also counted the numbers of GSCs in each of the imaged germaria and compared how many germaria contained one, two, or three GSCs. While Ameku and Niwa (2016) reported an increased number of germaria with three GSCs in mated females, we did not observe a significant difference in the numbers of germaria with one, two, or three GSCs among non-mated and mated females (Figure 2D).

One explanation for the different results could be that we performed single-female mating experiments while Ameku and Niwa (2016) used a mass-mating protocol. Therefore, we repeated the experiments under the same conditions as reported by Ameku and Niwa (2016). However, we obtained the same results with the mass-mated females as we did for single-mated females. No increase in MI^GSC^ was apparent (Figure 2E) and no increase in the numbers of germaria with three GSCs was seen (Figure 2F). We conclude that, in our hands, mating had no effect on female MI^GSC^ or GSC numbers.

While we didn’t detect differences in MI^GSC^ between virgin and mated females, we did observe the expected differences in oviposition. Previous studies have shown that virgin females tend to retain their eggs compared to mated females (Bouletreau-Merle et al., 1982; Bouletreau-Merle, 1990). Consistent with this, we observed that virgin *OR* females laid far less eggs compared to mated *OR* females (Table 2). The majority of the virgin females had not laid any eggs by 24 (176/193=91%) and 48 (117/144=81%) hours of the experiment, and the remainder of the females had laid 10 eggs by 24 hours and 20 eggs by 48 hours (Figure 2G, 2H). In comparison, a significantly smaller number of mated females had not laid eggs by either timepoint (44/204=22% and 7/132=0.5%) and most of the mated females had laid up to 20 eggs by 24 hours and up to 50 eggs by 48 hours of the experiment (Figure 2G, 2H). In accordance with having received sperm, 90% of the eggs laid by mated females (n>500) developed into larvae. As expected, eggs laid by the virgin females did not develop into larvae (n=70).

**Table 2.** Egg production. Eggs laid by virgin and mated *OR* females were counted and grouped into units of 10, n= number of females.

|  |  | # of eggs |  |  |  |  |  |  |
| --- | --- | --- | --- | --- | --- | --- | --- | --- |
| Condition | n | 0 | 1-10 | 11-20 | 21-30 | 31-40 | 41-50 | >50 |
| Virgin, 24 hours | 193 | 176 | 16 | 1 |  |  |  |  |
| Mated, 24 hours | 204 | 44 | 83 | 70 | 6 | 1 |  |  |
| Virgin, 48 hours | 144 | 117 | 23 | 1 | 1 |  |  |  |
| Mated 48 hours | 132 | 7 | 17 | 33 | 19 | 25 | 31 | 1 |

According to the literature, less eggs mature in ovaries of virgin compared to mated females, most likely due to the mid oogenesis checkpoint (Soller et al., 1997; Soller et al., 1999). After a female GSC daughter has undergone transit amplifying mitotic divisions, one of the 16 germline cells will become the future oocyte (blue in Figure 2A) and the other 15 germline cells will become nurse cells. By this time, the cluster of germline cells has reached region two of the germarium, where it becomes surrounded by somatic follicle cells to form an egg chamber. Once egg chambers have left the germarium, they develop through fourteen stages of oogenesis into mature eggs. During egg chamber maturation, the nurse cells provide the future oocyte with mRNAs, proteins, and nutrients while the follicle cells set the embryo axes and produce the vitelline membrane and the chorion that surround the mature egg (King, 1970; Spradling, 1993; Wu et al., 2008). The oocyte becomes morphologically distinct from the nurse cells at stage eight, when it starts to grow in size. Shortly after this, around stage nine, the mid oogenesis checkpoint occurs. Most likely, the checkpoint prevents the flies from investing too much energy into the development of mature eggs if conditions for reproduction are not favorable. The further development of the egg chambers is influenced by various factors, including diet, temperature, and mating (McCall, 2004; Pritchett et al, 2009). Specifically, the endocrine hormone, 20-hydroxy-ecdyson (20HE), promotes the progression through further egg chamber development. Animals without 20HE, or animals lacking the receptor or its downstream transcription factors accumulate degenerating egg chambers (Buszczak et al., 1999).

Interestingly, this stage is also regulated by SP, one of the key hormones produced in the accessory glands of the male reproductive tract and transferred to the female with the ejaculate. SP promotes yolk production specifically starting at stage nine of oogenesis, but also mediates several other post-mating responses, including female receptivity, oviposition, and ovulation (Chen et al., 1988; Soller et al., 1997; Chapman et al., 2003; Liu and Kubli, 2003). Recent advances have shown that SP promotes ovulation through a distinct neural circuit (Wang et al., 2020).

Our findings on female oviposition and fertility upon mating are consistent with the literature and with the previous report by Ameku and Niwa (2016). It is not clear to us why we did not detect the reported increase in MI^GSC^ and GSC numbers in mated females. It could be due to fluctuations in the quality of the molecular tools, specifically the antibodies, or differences in human analysis and interpretation of the results. In addition, or alternatively, one factor influencing the different outcomes may be the number of germaria analyzed. To spread our results over as many females as possible, we analyzed a limited number of germaria per ovary, while it appears that Ameku and Niwa (2016) analyzed every germarium from only a few ovaries. Finally, it is possible that our fly strains behave differently due to intrinsic parameters, or that air pressure or other environmental factors in the different countries have impacted the results.

## Methods

### Fly husbandry

Flies were raised on a cornmeal/agar diet and kept in incubators with temperature-, light-, and humidity-control. The *OR*, *CS*, and *w^1118^* flies were originally obtained from the Bloomington stock center but have been in the laboratory for more than 10 years, while the *w^1^* (BL#679) flies were obtained more recently (The Flybase Consortium, 2003). All four fly lines were repeatedly isogenized. *X*⌃*X*, *y, w, f* / Y / *shi^ts^* flies were obtained from Barry Ganetzky.

### Mating experiments

Mating experiments were performed at 29°C. Flies were fed with yeast paste on apple juice-agar plates in egg lay containers for 24 hours prior to the experiment. Male mating: Males were placed into one slot of a mating chamber either by themselves (non-mated) or with three virgin females (mated). The chambers were closed by apple juice-agar lids supplemented with yeast paste. On each of the following two days, females were discarded, and each mated male was provided with three new virgin females. Apple juice-agar lids supplemented with yeast paste were replaced daily for both conditions. Except for the fertility testes, females from the stock *X*⌃*X*, *y, w, f* / Y / *shi^ts^* were used for male mating experiments.

Female mating: For single female mating, three-day old virgin females were placed either by themselves or together with two one-week old males into one slot of the mating box. The chambers were closed with apple juice-agar lids supplemented with yeast paste. For mass mating, we prepared a total of 15 vials supplemented with yeast paste. Five of the vials were filled with 45 three-day old virgins each and 10 of the vials were filled with 15 three-day old female virgins and 30 one-week old males. After six hours, males were removed, and the females were dissected 24 hours later.

### Fertility tests and Oviposition

For male fertility tests, 100 females mated to males on day one of the experiment were placed into one food vial each and the vials were inspected for progeny after one week. For female fertility, the flies were mated in vials or mating boxes for six hours as described above, and the vials or apple juice/agar plates were inspected for progeny after 72 hours. Eggs were counted by eye on the apple juice/agar plates from non-mated and mated *OR* females after 24 and 48 hours under a dissection microscope.

### Immuno-fluorescence and microscopy

Animals were placed on ice to immobilize them. Gonads were dissected in Tissue Isolation Buffer (TIB) and immediately placed into a 1.5 ml tube with cold TIB buffer. Gonads were then fixed, stained and imaged as previously described (Schulz et al., 2002; Parrott et al., 2012). The mouse anti-FasciclinIII (FasIII) antibody (1:10) developed by C. Goodman and the mouse anti-LaminC antibody (1:10) developed by P. A. Fisher were obtained from the Developmental Studies Hybridoma Bank, created by the NICHD of the NIH and maintained at The University of Iowa, Department of Biology, Iowa City, IA 52242. The goat anti-Vasa antibody (1:50 to 1:500) was purchased from Santa Cruz Biotechnology Inc. (sc26877). The rabbit anti-phosphorylated Histone H3 (pHH3) antibodies (1:100 to 1:1000) were purchased from Fisher (PA5-17869), Millipore (06-570), and Santa Cruz Biotechnology Inc. (sc8656-R). Secondary antibodies coupled to Alexa 488, 568, and 647 (1:1000) and Slow Fade Gold/DAPI embedding medium were purchased from Life Technologies. Staining was observed with a Zeiss Axiophot, and images taken with a digital camera and an apotome via the Axiovision Rel. software.

### Graphic presentations and data analysis

Bar graphs and FDGs were generated with GraphPad prism version 7 and assembled into composite images using Adobe Photoshop. The GraphPad prism default two-tailed student’s t-test was used to analyze statistical relevance.

## ACKNOWLEDGEMENTS

The authors are grateful to Leon McSwain, Alicia Hudson, and Kyona Garrett for technical assistance. We thank Bruce Baker for helpful discussions, and Heath Aston for art work and critical comments on the manuscript. We are grateful to Barry Ganetzky for the *X*⌃*X*, *shi^ts^* fly stock and to Wolfgang Lukowitz for the use of his microscope. This work was supported by NSF grants #0841419 and #1355009.

## Author contributions

C.S developed and supervised the project, and M. M. and C.S. performed the experiments and wrote the manuscript.

## Competing interests

The authors declare no competing interests.

## Notes

### Competing Interest Statement

The authors have declared no competing interest.

